# YAP/TAZ Maintain the Proliferative Capacity and Structural Organization of Radial Glial Cells During Brain Development

**DOI:** 10.1101/2021.05.05.442840

**Authors:** Alfonso Lavado, Ruchika Gangwar, Joshua Paré, Shibiao Wan, Yiping Fan, Xinwei Cao

## Abstract

The Hippo pathway regulates the development and homeostasis of many tissues and in many species. It controls the activity of two paralogous transcriptional coactivators, YAP and TAZ (YAP/TAZ). Although previous studies have established that aberrant YAP/TAZ activation is detrimental to mammalian brain development, whether and how endogenous levels of YAP/TAZ activity regulate brain development remain unclear. Here, we show that during mammalian cortical development, YAP/TAZ are specifically expressed in apical neural progenitor cells known as radial glial cells (RGCs). The subcellular localization of YAP/TAZ undergoes dynamic changes as corticogenesis proceeds. YAP/TAZ are required for maintaining the proliferative potential and structural organization of RGCs, and their ablation during cortical development reduces the numbers of cortical projection neurons and causes the loss of ependymal cells, resulting in hydrocephaly. Transcriptomic analysis using sorted RGCs reveals gene expression changes in YAP/TAZ-depleted cells that correlate with mutant phenotypes. Thus, our study has uncovered essential functions of YAP/TAZ during mammalian brain development and revealed the transcriptional mechanism of their action.

## Introduction

All neurons and macroglia (astrocytes and oligodendrocytes) in the mammalian central nervous system (CNS) are derived from apical neural progenitor cells, which are also referred to as neuroepithelial cells and radial glial cells (RGCs) [reviewed in (Kriegstein and Alvarez-Buylla, 2009; Taverna et al., 2014; Villalba et al., 2021)]. These cells are organized as an epithelial sheet, with their cell bodies lining the lumen of the developing neural tube and forming the region known as the ventricular zone (VZ), their neighboring apical endfeet connecting with each other through tight junctions or adherens junctions at the ventricular surface, and their basal radial processes touching the pial basement membrane. At the early stages of cortical development (e.g., before embryonic days [E] 11–12 in mice), RGCs predominantly undergo symmetric proliferative division to expand the progenitor pool. They then mostly shift to asymmetric neurogenic division to self-renew and simultaneously produce neurons either directly or indirectly via intermediate progenitor cells (IPCs), which divide in the subventricular zone (SVZ) to produce more neurons. Towards the late stages of cortical development, RGCs mostly undergo terminal neurogenic division to exit the cell cycle, with a small fraction converting to glial progenitor cells and ependymal cells. Newborn neurons migrate outward through the intermediate zone (IZ) and settle at appropriate locations in the cortical plate (CP) to complete their differentiation, with early-born neurons occupying deep cortical layers and late-born neurons occupying upper layers.

The properties and behavior of neural progenitor cells, such as their proliferation, differentiation, metabolism, and structural organization, are intricately regulated to ensure the generation of a nervous system composed of the correct types and numbers of cells that are properly organized and connected. One node of this regulation is the Hippo pathway. Through complex signaling cascades, the Hippo pathway ultimately controls the activity of two paralogous transcriptional coactivators called YAP and TAZ (YAP/TAZ). When activated, YAP/TAZ enter the nucleus and interact with DNA-binding factors, mainly the TEAD factors, to regulate gene expression, often leading to increased cell proliferation and tissue growth or even tumorigenesis [reviewed in (Zheng and Pan, 2019)]. YAP/TAZ activity is tightly controlled during murine cortical development. When their activity is elevated by the loss of the neurofibromatosis type 2 (NF2/Merlin) tumor suppressor, neural progenitors overexpand and the differentiation of some neuronal and glial cell types is perturbed, resulting in impaired morphogenesis of the hippocampus and agenesis of the corpus callosum (Lavado et al., 2013; Lavado et al., 2014). The loss of partitioning defective 3 (PARD3), a protein involved in asymmetric cell division, upregulates YAP/TAZ activity during early stages of cortical development, resulting in ectopic RGC production and the formation of abnormal neuron clusters known as heterotopia (Liu et al., 2018). Removal of centrosomal protein 83 (CEP83) also activates YAP and causes excessive proliferation of RGCs, which leads to the formation of an enlarged cortex with abnormal folding (Shao et al., 2020). Most strikingly, when YAP/TAZ activity is unleashed by the inactivation of LATS1 and LATS2—the direct inhibitors of YAP/TAZ and the gatekeepers of YAP/TAZ activity—neural progenitor cells over-proliferate transiently and quickly undergo massive cell death due to the replication stress triggered by global hypertranscription (Lavado et al., 2018). Although these findings demonstrate the importance of restraining YAP/TAZ activity to enable normal cortical development, the role of endogenous YAP/TAZ activity in cortical RGCs has not been analyzed carefully. In *Yap* conditional knockout (cKO) mice in which *Yap* is deleted by the *Nestin-Cre* line, which expresses the Cre recombinase in neural progenitor cells throughout the CNS starting at approximately E10.5 (Tronche et al., 1999), the proliferation and number of cortical RGCs are not affected (Huang et al., 2016; Park et al., 2016). Instead, astrocyte proliferation is reduced (Huang et al., 2016) and the generation of ependymal cells lining the aqueduct is impaired (Park et al., 2016). However, TAZ function remains intact in these *Yap* cKO mice and may compensate for the loss of YAP in cortical RGCs.

In this study, we found that deleting both *Yap* and *Taz* in murine cortical RGCs severely perturbed cortical development. YAP/TAZ loss slowed the cell-cycle speed of RGCs and caused premature cell-cycle exit, resulting in reduced numbers of cortical projection neurons. YAP/TAZ loss also impaired the structural organization of the progenitor zone and caused loss of ependymal cells. Our RNA-sequencing analysis of sorted RGCs revealed gene expression changes in YAP/TAZ-depleted RGCs that correlated with mutant phenotypes, suggesting that YAP/TAZ act by regulating gene expression.

## Materials and Methods

### Mice

*Yap^F/F^;Taz^F/F^* mice were obtained from Dr. Eric Olson (University of Texas Southwestern Medical Center at Dallas) (Xin et al., 2011). *Emx1-Cre* mice were obtained from The Jackson Laboratory (stock #005628). All mice were maintained in a mixed genetic background. All animal procedures were approved by the Institutional Animal Care and Use Committee of St. Jude Children’s Research Hospital. Mouse housing and husbandry conditions followed the standards set by the Animal Resource Center at St. Jude Children’s Research Hospital.

### Western blot analysis

E14.5 mouse cortices were dissected, and total cell lysates were prepared in 20 mM HEPES (pH 7.4), 150 mM NaCl, 2% SDS, and 5% glycerol supplemented with AEBSF (Sigma-Aldrich A8456) and Halt protease and phosphatase inhibitors (Thermo Fisher Scientific #78440). Protein concentrations were measured by BCA assay (Thermo #23225). 20 μg of protein per lane were subjected to SDS-PAGE and probed with the following primary antibodies: YAP (Cell Signaling Technologies #8418, 1:500), YAP/TAZ (Cell Signaling #4912, 1:500), and GAPDH (Cell Signaling #2118, 1:5000).

### Histology, immunostaining, and TUNEL assay

Embryonic mouse brains were dissected in PBS and fixed overnight in 4% paraformaldehyde (PFA) at 4°C. Postnatal animals were perfused with 0.9% saline and PFA, and their brains were dissected and fixed in PFA overnight at 4°C. Histology, immunostaining, and TUNEL assay were performed as described previously (Lavado et al., 2018). Luxol blue and cresyl violet staining was performed using the Kluver-Barrera method (EMS #26681). The primary antibodies used were: YAP/TAZ (Cell Signaling #8418, 1:500), SOX2 (Santa Cruz Biotechnology #SC17320, 1:100), SOX2–Alexa-647 (BD Biosciences #562139, 1:100), TBR2 (Thermo #14-4875-82, 1:250), TBR2 (Abcam ab23345, 1:250), TBR1 (Abcam ab31940, 1:1000), CUX1 (Santa Cruz #SC13024, 1:250), CTIP2 (Abcam ab18465, 1:1000), PAX6 (BioLegend #901301, 1:500), BrdU/IdU (BD Biosciences #347580, 1:50), BrdU (Abcam ab152095, 1:500), Ki67 (Vector laboratories #VP-RM04, 1:500), ZO1– Alexa-488 (Thermo #339188, 1:100), Nestin (R&D systems #AF2736, 1:100), ARL13b (ProteinTech #17711-1-AP, 1:250), β-Catenin–Alexa-647 (Cell Signaling #4627, 1:100), and S100β (Sigma #52532, 1:100).

### Image acquisition and quantification

Bright-field images were acquired using a Zeiss Stereo Discovery V8 microscope equipped with an AxioCam MRc camera and were processed with AxioVision software (Zeiss). Fluorescence images were acquired using a Zeiss LSM 780 confocal microscope. All images used for cell counting were acquired using a Marianas spinning disk confocal microscope or a Zeiss LSM 780 confocal microscope with a 40× objective. Cells were counted using Imaris 9.1 (Bitplane) or the machine learning–based image segmentation method StarDist (Schmidt et al., 2018). In the latter case, StarDist was used to detect individual cells in multi-channel images and the average fluorescence intensity of each cell was quantified for each channel. The detection and quantification steps were implemented as a batch execution script in the OMERO server platform (Allan et al., 2012). The OMERO viewer was used to interactively select the intensity threshold in each channel for cells positive for the marker. For each genotype, 3–6 embryos with 1–5 sections per embryo at comparable positions were quantified. For quantifying cell death, TUNEL positive cells were counted manually. For each genotype, 4–6 embryos with 3 sections of each rostrocaudal position per embryo were quantified. Data points in all quantification graphs correspond to individual embryos.

### Quantification of cell-cycle exit and cell-cycle length

To measure cell-cycle exit at E14.5 and E16.5, EdU (10 μg/g body weight) was injected intraperitoneally into pregnant dams at E13.5 and E15.5, respectively, and embryos were collected 24 h later. EdU was detected by the Click-it EdU Alexa Fluor 647 Imaging Kit (Thermo Invitrogen) after immunostaining with a Ki67 antibody. The cell-cycle exit index was calculated as the fraction of EdU+ cells that were Ki67− (i.e., EdU+Ki67−/EdU+) and was represented as a percentage. For cell-cycle exit quantification at E17.5, pregnant dams were injected intraperitoneally with IdU (50 μg/g body weight) at E16.5. Cell-cycle lengths were measured as described previously (Lavado et al., 2018).

### MARIS (method for analyzing RNA following intracellular sorting)

MARIS was performed using a modified protocol from (Baizabal et al., 2018). Neocortices from E16.5 embryos were dissected and dissociated into single cell suspensions by using StemPro Accutase (Thermo #A1110501) for 3 min at 37°C. The following steps were performed at 4°C. Cells were collected by centrifugation, gently resuspended in fixation buffer (PBS, 0.1% saponin, 4% PFA, RNAsin [New England Biolabs #M0314] 1:20) and incubated for 30 min. Cells were collected by centrifugation and washed twice with wash buffer (PBS, 0.1% saponin, 0.1% BSA, RNAsin 1:100) then incubated with the primary antibodies, rabbit anti-PAX6 (BioLegend #901301, 1:2000) and mouse anti-TBR2-APC (MACS #130-102-378, 1:25), in antibody buffer (PBS, 0.1% saponin, 1% BSA, RNAsin 1:20) for 1 h with gentle rocking. Cells were then washed twice with wash buffer and incubated with an anti-rabbit Alexa Fluor 488–conjugated secondary antibody (Jackson ImmunoResearch Laboratories) in antibody buffer for 30 min. After two washes with wash buffer, cells from two *Yap^F/F^;Taz^F/F^* control brains or three *Yap^F/F^;Taz^F/F^;Emx-Cre* dKO brains were resuspended in 1 ml of sorting buffer (PBS, 0.5% BSA, RNAsin 1:20) and filtered through a 40-μm Flowmi cell strainer (Sigma #136800040). Cells were sorted on a FACSAria cell sorter (BD Biosciences) using a 100-μm nozzle. The PAX6+TBR2− and PAX6−TBR2+ fractions were collected in tubes containing PBS, 0.5% BSA, and RNAsin 1:4. After sorting, cells were collected by centrifugation at 3000 × *g* for 10 min. Total RNA was isolated using the RecoverAll Total Nucleic Acid Isolation Kit (Thermo #AM1975) and concentrated using the RNeasy MinElute Cleanup Kit (Qiagen #74204). Library preparation and sequencing were performed by the Genome Sequencing Facility at St. Jude Children’s Research Hospital. RNA quality was checked by a 2100 Bioanalyzer RNA 6000 Pico Assay (Agilent) before library generation. Libraries were prepared using the Ovation RNA Seq V2 Kit (NuGEN) and quantified using the Quant-iT PicoGreen dsDNA Assay (Life Technologies). Samples were sequenced with an Illumina HiSeq4000 system, yielding 100 million 100-bp paired-end reads. Five control and four dKO libraries were sequenced.

### RNA-seq data analysis

Sequences were mapped to the mm10 genome by using the STAR aligner (Dobin et al., 2013). Transcript-level data was counted using HTSEQ (Anders et al., 2015). Differential gene expression was modeled using voom (Law et al., 2014), available in the limma R software package. Normalization factors were generated using the TMM method. The voom-normalized counts were analyzed using the lmFit and eBayes functions in limma. The adjusted *P* value was estimated using the Benjamini-Hochberg method. Gene set enrichment analysis (GSEA) was performed using gene sets obtained from MSigDB (the Broad Institute) as described previously (Subramanian et al., 2005). The ranking of genes was calculated using the signal-to-noise algorithm, and a *P*-value for each gene set was estimated by comparing the observed enrichment score to that obtained from a null distribution computed from 1000 permutations of genes within the gene sets. The false-discovery rate (FDR) was estimated as described previously (Subramanian et al., 2005).

## Results

### YAP/TAZ exhibit dynamic subcellular localization in RGCs during cortical development

To study the role of YAP/TAZ during cortical development, we first examined their expression and localization by immunostaining using an antibody that recognizes both YAP and TAZ. At E12.5, YAP/TAZ immunoreactivity was highly enriched at the ventricular surface, with weak cytoplasmic staining also being detected in the VZ demarcated by the RGC marker SOX2 (Figure 1A). At E14.5, in addition to their strong expression at the ventricular surface, YAP/TAZ were detected diffusely in the cytoplasm and nucleus of RGCs (Figure 1B). Interestingly, at E16.5, YAP/TAZ became enriched in the nucleus of RGCs while still being present at the ventricular surface (Figure 1C). Therefore, YAP/TAZ proteins shift from the cytoplasm to the nucleus in RGCs as cortical development proceeds.

**Figure 1.**
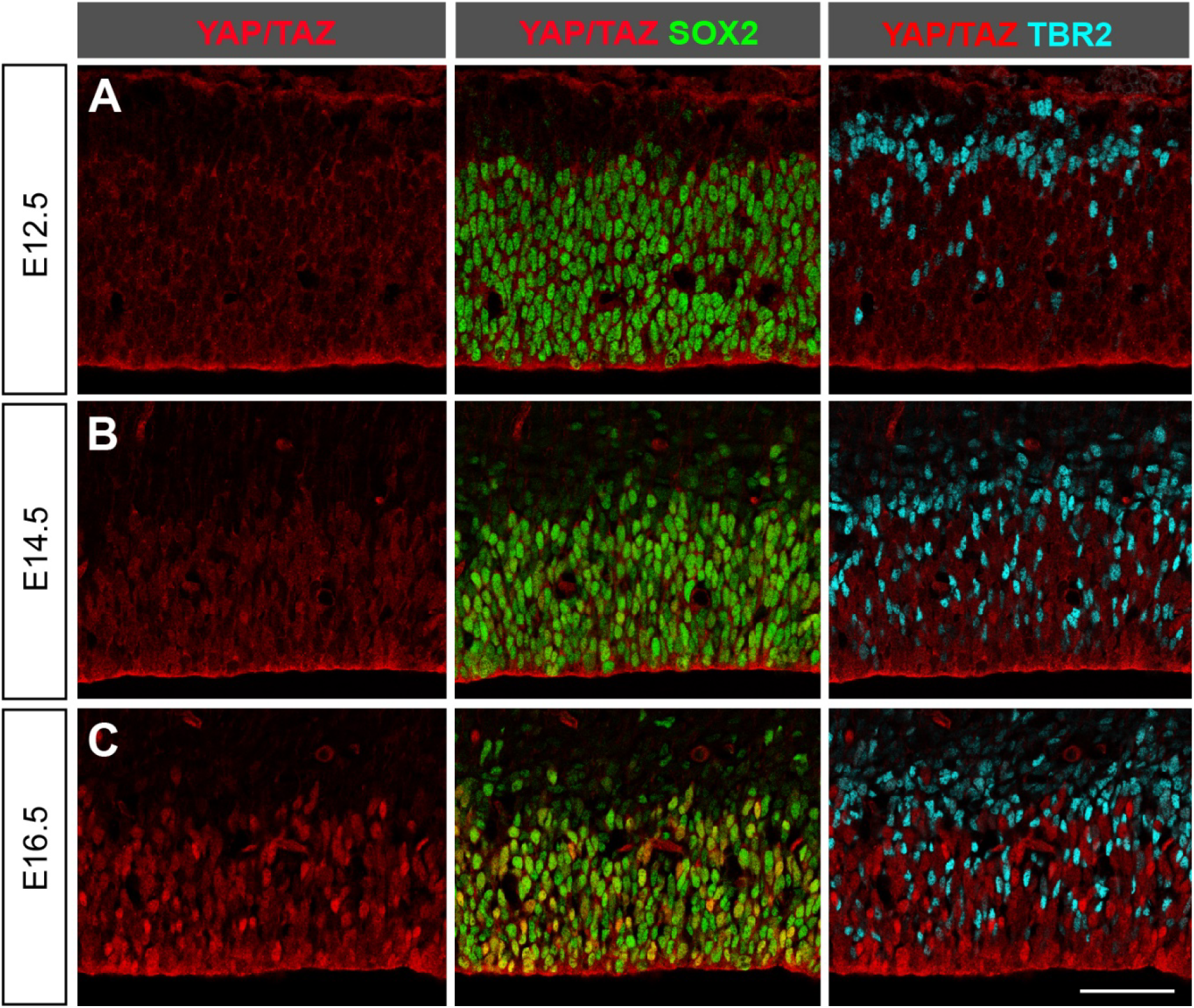
YAP/TAZ exhibit dynamic subcellular localization in RGCs during cortical development. (**A**–**C**) Immunostaining with antibodies against YAP/TAZ, the radial glial cell (RGC) marker SOX2, and the intermediate progenitor cell (IPC) marker TBR2 during murine cortical development. Scale bar, 50 μm.

To determine whether YAP/TAZ were also expressed in IPCs, we performed co-staining with an antibody against the IPC marker TBR2 (Englund et al., 2005). No YAP/TAZ immunoreactivity was detected in TBR2+ IPCs at any developmental stage (Figure 1A–C). Therefore, YAP/TAZ are specifically expressed in RGCs but not in IPCs.

### The loss of YAP/TAZ causes severe brain malformation

To determine the role of YAP/TAZ during cortical development, we generated *Yap* (*Yap1*) and *Taz* (*Wwtr1*) cKO mice by crossing *Yap^F/F^;Taz^F/F^* mice to the *Emx1-Cre* line, which expresses Cre selectively in the developing cortex starting at approximately E9.5 (Gorski et al., 2002). Immunostaining showed that YAP/TAZ signals were depleted in the cortices of *Yap^F/F^;Taz^F/F^;Emx1-Cre* (referred to as *Yap;Taz* double-knockout [dKO]) embryos at E12.5 (Figure 2A). Western blot analysis of E14.5 cortices showed a marked reduction in YAP and TAZ proteins in the dKO cortices as compared to the control (no Cre) cortices (Figure 2B).

*Yap;Taz* dKO mice were born at the expected Mendelian ratio and often survived past 1 month of age. However, the brains of 1-month-old dKO mice exhibited severe abnormalities. Their cortices were much thinner than control cortices, and their lateral ventricles were drastically enlarged (Figure 2C), a condition known as hydrocephaly. Mice lacking three of the four alleles of the *Yap;Taz* genes occasionally exhibited abnormalities. To compare the severity of brain malformation in different genotypes, we devised a simple scoring system based on the most recognizable defect, the enlargement of the lateral ventricles. Brains with no visible enlargement were assigned a score of 0, those with an enlargement in a restricted region of just one lateral ventricle a score of 1, and those with enlargements throughout the anterior–posterior axis of one or both lateral ventricles a score of 2 or 3, respectively (Figure 2D). All control and *Yap^F/+^;Taz^F/+^;Emx1-Cre* mice had a score of 0 as they exhibited no obvious defects. The vast majority of mice lacking three of the four *Yap;Taz* alleles also had a score of 0. In contrast, all *Yap;Taz* dKO brains had a score of 2 or 3 (Figure 2E). These results indicate that YAP/TAZ are required for normal brain development and that they function in a largely redundant manner.

**Figure 2.**
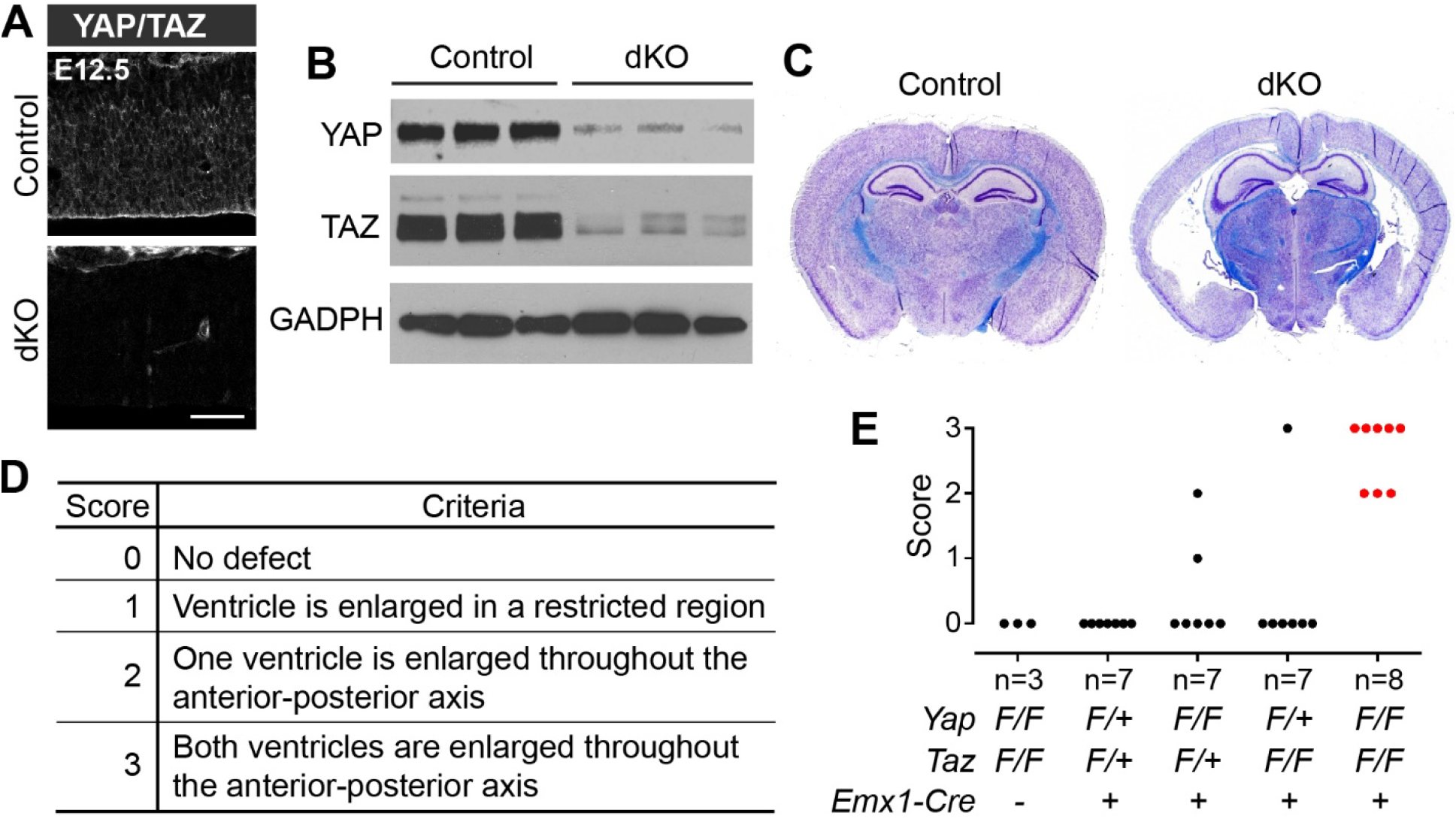
YAP/TAZ depletion causes brain malformation. (**A**) Immunostaining showing loss of YAP/TAZ signals in *Yap;Taz* dKO cortex at E12.5. Scale bar, 50 μm. (**B**) Western blot analysis showing reduced YAP and TAZ protein levels in dKO cortex at E14.5. (**C**) Nissl and Luxol blue staining showing gross morphologic abnormalities in dKO brains at postnatal day (P) 30. (**D**) The scoring system used for quantifying brain defects. (**E**) Quantification of the brain phenotype severity according to the genotype.

### Loss of YAP/TAZ reduces the number of neurons

To understand the origins of brain malformation caused by *Yap;Taz* deletion, we examined the cortices of *Yap;Taz* dKO mice at several developmental stages. We focused on the neocortex for all our analyses. First, we analyzed cortical projection neurons, which are generated from RGCs and IPCs in the VZ and SVZ and migrate into the cortical plate. The position occupied by these neurons depends on when they are born, with deep-layer (V, VI) neurons being born at E12.5 to E13.5 and up-layer (II–IV) neurons being born from E14.5 onwards [reviewed in (Greig et al., 2013; Kwan et al., 2012)]. The number of early-born TBR1+ layer VI neurons was unchanged in the *Yap;Taz* dKO cortex when compared to that in the control cortex at E14.5 but was significantly reduced at E16.5 and E17.5 (Figure 3A–D). To quantify later-born up-layer neurons, we restricted our analysis to postnatal day (P) 0 because, at that stage, the cortical thickness and lateral ventricle size were similar in control and *Yap;Taz* dKO brains (data not shown). The numbers of CTIP2+ layer V neurons (dashed lines, Figure 3E, G) and CUX1+ layers II–IV neurons (dashed lines, Figure 3F, H) were significantly reduced in the *Yap;Taz* dKO cortex. The positioning of neurons of different layers was largely normal in the *Yap;Taz* dKO cortex (Figure 3C, E, F). Together, these results show that the lack of YAP/TAZ reduces the numbers of both early-born and late-born cortical projection neurons.

**Figure 3.**
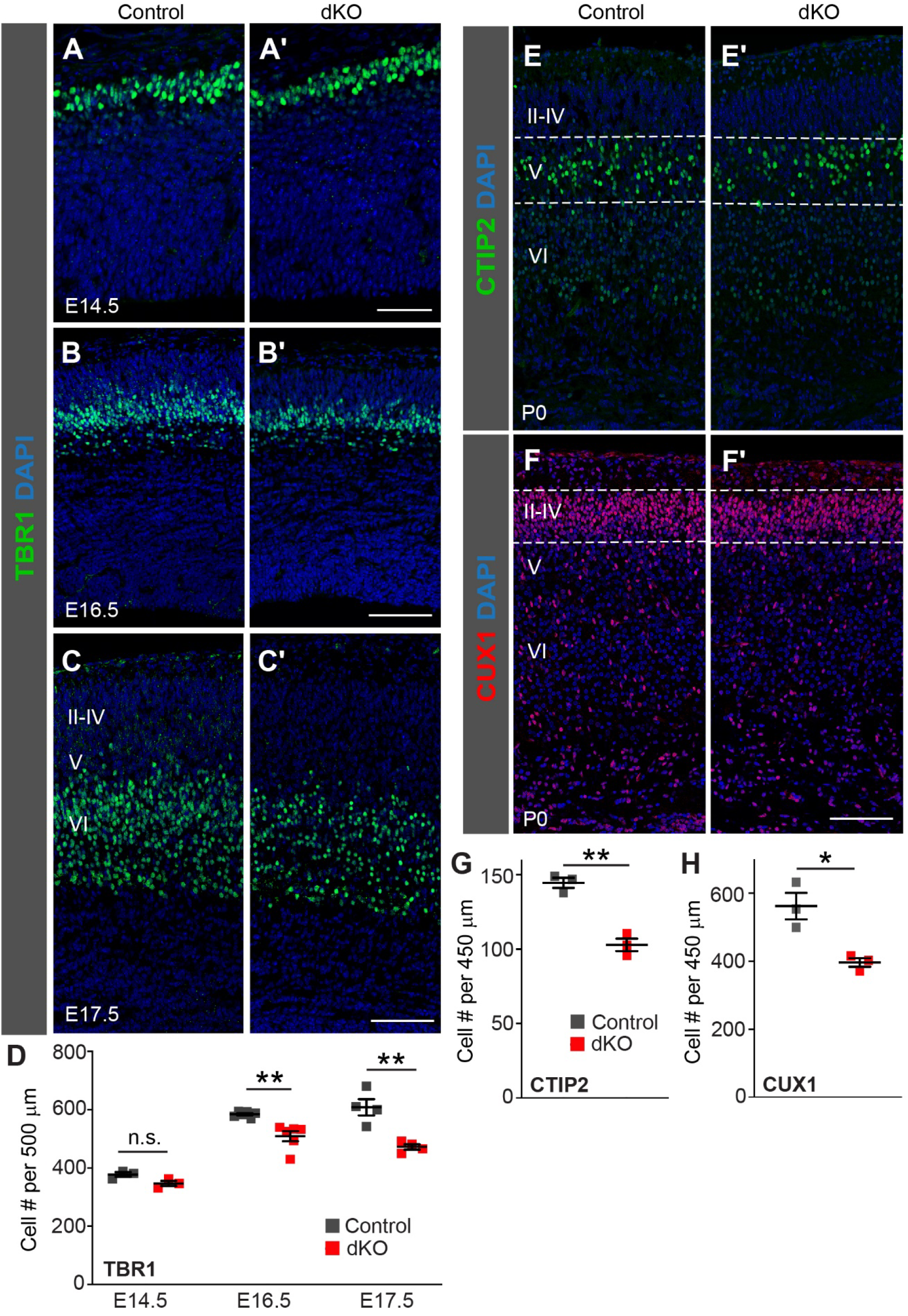
Loss of YAP/TAZ reduces the number of neocortical neurons. (**A**–**D**) Immunostaining and quantification of TBR1+ layer VI neurons in control and *Yap;Taz* dKO cortices. (**E–H**) Immunostaining and quantification of CUX1+ layer II–IV neurons and CTIP2+ layer V neurons in P0 cortices. Dashed lines indicate the regions quantified. Data are shown as mean ± SEM; * *P* < 0.05; ** *P* < 0.01; n.s., not significant; unpaired two-tailed Student’s *t* test. Scale bars, 50 μm in A, 100 μm in B, C, E, F.

### YAP/TAZ maintain the proliferative capacity of cortical RGCs

To uncover how YAP/TAZ depletion in RGCs led to decreased numbers of cortical neurons, we first examined the effect of *Yap;Taz* deletion on progenitor cell populations during cortical development. At E14.5, the numbers of SOX2+TBR2− RGCs and TBR2+ IPCs were similar in control and *Yap;Taz* dKO cortices (Figure 4A, D, E). However, at E16.5 and E17.5, the numbers of both RGCs and IPCs were significantly reduced in *Yap;Taz* dKO cortices as compared to control cortices (Figure 4B–E). In addition, the organization of the VZ/SVZ, the region densely populated by SOX2+ and TBR2+ progenitor cells in the normal developing cortex (Figure 4B, C), was notably disrupted in some regions of the dKO cortex starting at E16.5. In these defective regions, both RGCs and IPCs appeared to be more dispersed in the dKO cortex, with many progenitor cells being mispositioned in the IZ, which is normally occupied by newborn neurons (Figure 4B’). Quantification showed that the number of RGCs in the IZ was significantly increased in the dKO cortex at E16.5 (Figure 4G). At E17.5, the progenitor-dense VZ/SVZ was no longer discernable in some regions of the dKO cortex (Figure 4C’). Therefore, the lack of YAP/TAZ reduces the numbers of progenitor cells in the neocortex and disrupts the organization of the VZ/SVZ.

**Figure 4.**
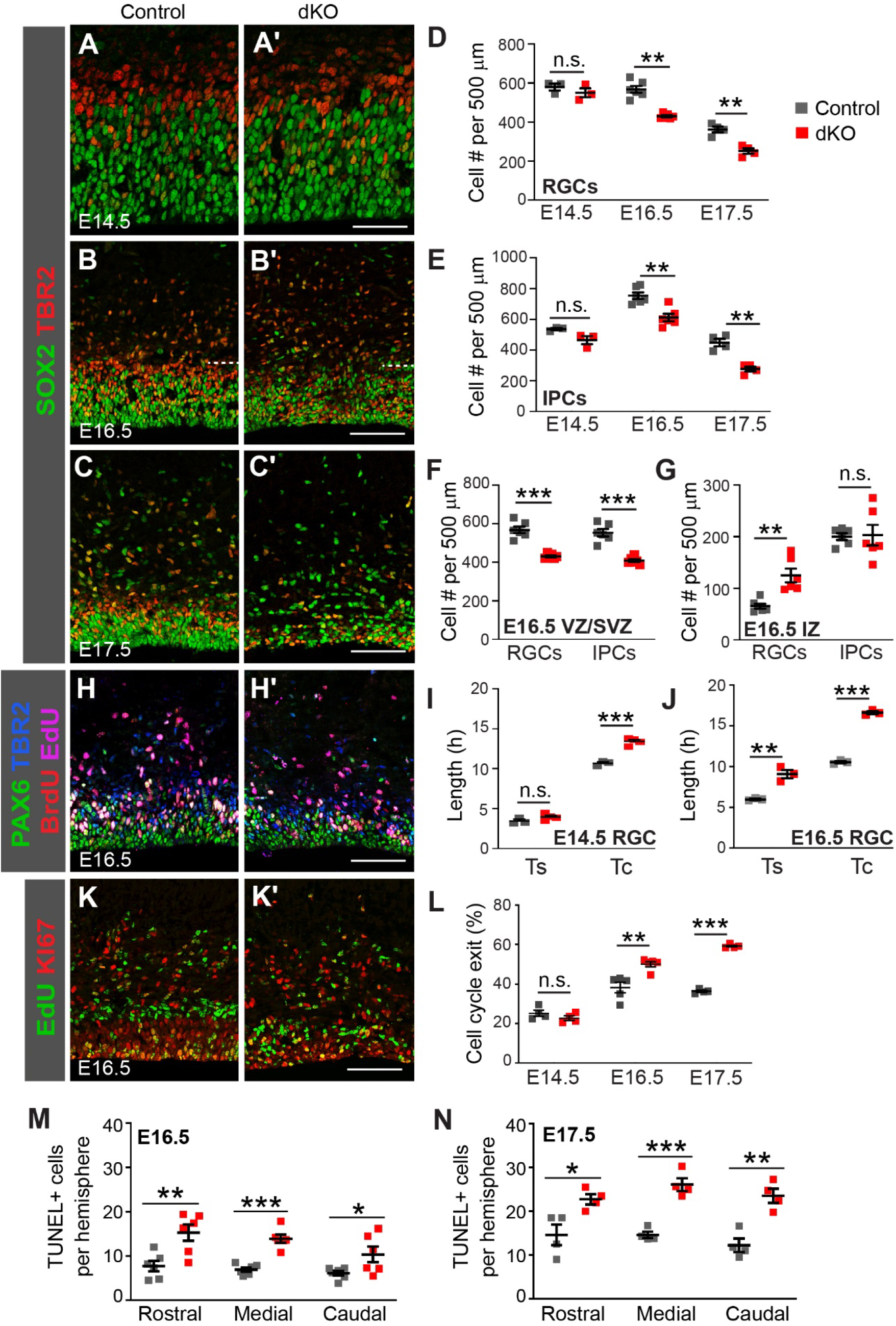
Loss of YAP/TAZ reduces the proliferative capacity of progenitor cells and increases cell death. (**A**–**C**) SOX2 and TBR2 immunostaining in control and *Yap;Taz* dKO cortices. Dashed line in B marks the boundary between the VZ/SVZ (ventricular zone/subventricular zone) and the IZ (intermediate zone). (**D**) Quantification of the numbers of neocortical SOX2+TBR2- RGCs. (**E**) Quantification of the numbers of neocortical TBR2+ IPCs. (**F**, **G**) Quantification of the numbers of RGCs and IPCs in the VZ/SVZ (F) and in the IZ (G) at E16.5. (**H**–**J**) Immunostaining and quantification of the S-phase length (Ts) and the total cell-cycle length (Tc) of PAX6+TBR2- RGCs by cumulative BrdU/EdU-labeling. Representative E16.5 immunostaining images are shown.(**K**, **L**) Immunostaining and quantification of cell-cycle exit. Representative E16.5 immunostaining images are shown. (**M**, **N**) Quantification of TUNEL assay signals in rostral, medial, and caudal levels of the neocortex. Data are shown as mean ± SEM; * *P* < 0.05; ** *P* < 0.01; *** *P* < 0.001; n.s., not significant; unpaired two-tailed Student’s *t* test. Scale bars, 50 μm in A, 100 μm in B, C, H, K.

To understand the cause of the reduction in progenitor cell numbers in the dKO cortex, we analyzed the proliferative capacity of RGCs. We measured the cell-cycle length of RGCs by using a cumulative BrdU/EdU-labeling paradigm (Pilaz et al., 2016) (Figure 4H). At E14.5, the total cell-cycle length (Tc) of dKO RGCs was significantly longer than that of control RGCs, although their S-phase lengths (Ts) were similar (Figure 4I). At E16.5, both the Tc and Ts of dKO RGCs were longer than those of control RGCs (Figure 4J). We also measured the cell-cycle exit of progenitor cells by using a 24-h EdU-labeling paradigm. The fraction of progenitor cells that exited the cell cycle at E14.5 was unchanged in the dKO cortex when compared to that in the control cortex but was increased at E16.5 and E17.5 (Figure 4K, L). Therefore, the loss of YAP/TAZ slows the cell-cycle speed of RGCs and causes premature cell-cycle exit, leading to a reduction in progenitor cell numbers and, consequently, a reduction in neuron numbers.

In addition to the proliferation defects, cell death was significantly but moderately elevated in the dKO cortex at E16.5 and E17.5 when compared to that in the control cortex (Figure 4M, N), which probably also contributed to the reduction in the number of cortical neurons.

### YAP/TAZ maintain the structural organization of cortical progenitor cells

Next, we analyzed in more detail the VZ/SVZ organizational defect observed in the *Yap;Taz* dKO cortex. In the normal developing cortex, SOX2+ RGCs are densely packed in the VZ and most TBR2+ IPCs reside above the RGCs in the SVZ (Figure 5A). RGCs are connected with each other through adherens junctions located at the ventricular surface (Figure 5A, C, E), which maintain the tissue integrity of the neuroepithelium. Each RGC harbors a single primary cilium at the apical surface that protrudes into the ventricle [reviewed in (Taverna et al., 2014)] (Figure 5E’, arrowheads). In some regions of the E16.5 *Yap;Taz* dKO cortex, SOX2+ RGCs were sparse and intermingled with TBR2+ IPCs, making it difficult to distinguish between the VZ and the SVZ (Figure 5B, bracket, B’), and TBR2+ IPCs were often located abnormally near the ventricular surface (Figure 5B’, arrowheads). In these defective regions, which in total encompassed approximately 30% of the cortical ventricular length, the adherens junction components ZO1 and β-catenin and the primary cilia labeled by ARL13B were completely lost from the ventricular surface (Figure 5B, D, F) and the radial glial processes labeled by Nestin were disorganized (Figure 5D), suggesting that the adherens junctions and apical-basal polarity of dKO RGCs were disrupted. Similar defects were observed in the E17.5 dKO neocortex, with the defective regions covering approximately 50% of the cortical ventricular length (data not shown). These analyses revealed that YAP/TAZ are required to maintain the structural organization and polarity of RGCs in the cortex.

**Figure 5.**
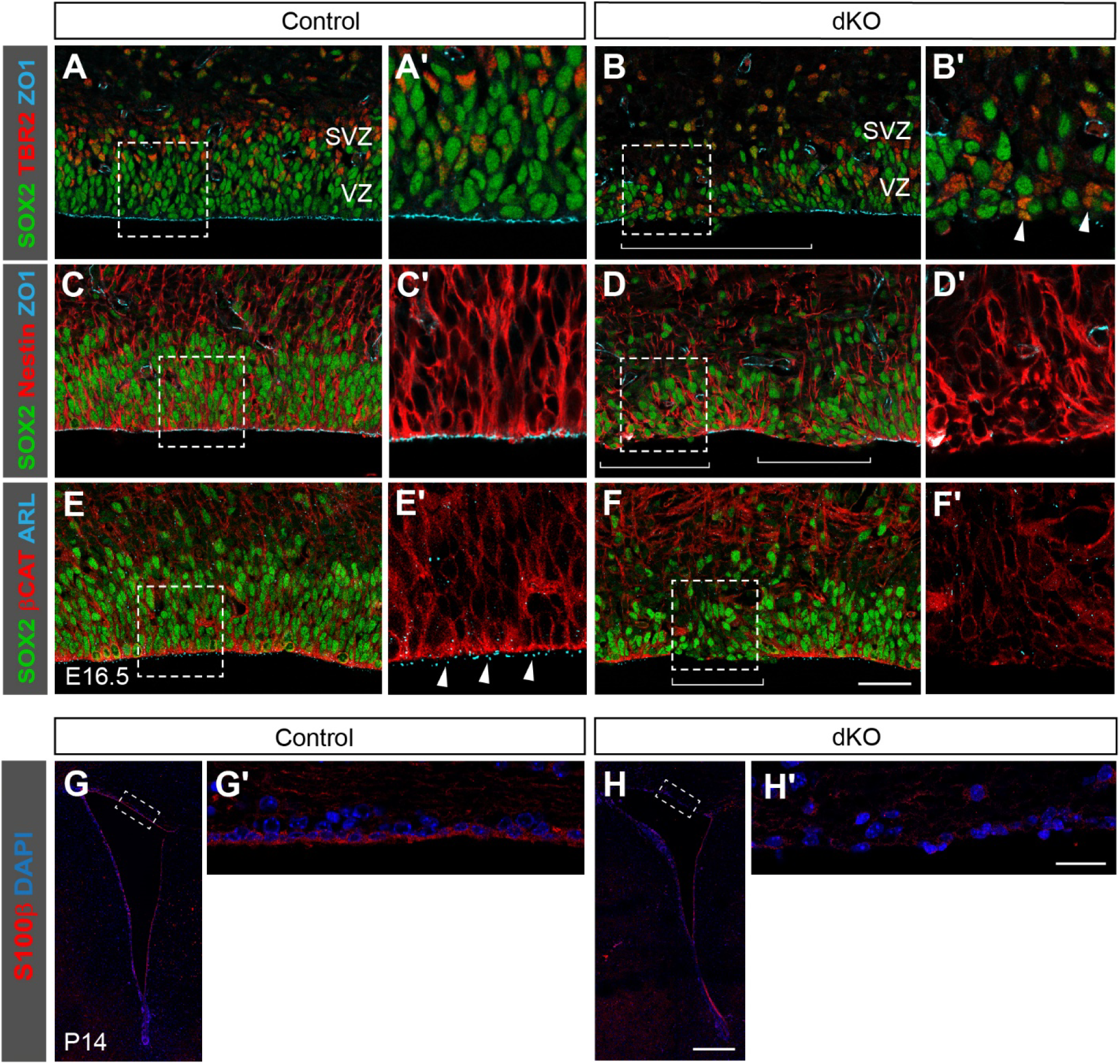
Loss of YAP/TAZ disrupts neuroepithelial organization and causes loss of ependymal cells. (**A**–**F**) Immunostaining with antibodies against the RGC marker SOX2, the IPC marker TBR2, the radial glial process marker Nestin, the adherens junction components ZO1 and β-catenin, and the primary cilium marker ARL13b, showing disorganization and disruption in the *Yap/Taz* dKO neocortex at E16.5. Brackets in B, D, and F mark areas with disruption. Arrowheads in B’ mark TBR2+ cells abnormally localized near the ventricular surface. Arrowheads in E’ mark primary cilia. (**G, H**) S100β immunostaining showing the lack of ependymal cells in the *Yap;Taz* dKO neocortex at P14. A’–H’ are enlargements of the boxed areas in A–H.

### YAP/TAZ depletion results in the loss of cortical ependymal cells

We next investigated the cause of hydrocephaly in *Yap;Taz* dKO brains. We focused on ependymal cells, which are multi-ciliated cells that line the walls of brain ventricles and control the flow of cerebrospinal fluid, because defects in ependymal cell function often result in hydrocephaly [reviewed in (Ohata and Alvarez-Buylla, 2016)] and because YAP is expressed in cortical ependymal cells (Eder et al., 2020) and is required for ependymal cell generation along the aqueduct (Park et al., 2016). Although ependymal cells are generated from RGCs during embryonic development (Spassky et al., 2005), their marker, S100β is not detectable in cortical ependymal cells by immunostaining until around P14. S100β+ ependymal cells, which lined the ventricular surface in the control brain (Figure 5G), were completely absent from the ventricular surface of the *Yap;Taz* dKO cortex (Figure 5H). This result showed that YAP/TAZ are required for the formation of the ependymal layer in the cortex, which probably underlies the hydrocephaly phenotype of *Yap;Taz* dKO brains.

### Transcriptional changes in RGCs caused by the loss of YAP/TAZ

To understand the molecular underpinning of the phenotypes of the *Yap;Taz* dKO cortex, we examined the impact of YAP/TAZ loss on RGC gene expression. We chose the E16.5 timepoint because RGC phenotypes first appeared in dKO cortices around this stage. We used the MARIS (method for analyzing RNA following intracellular sorting) method to immunolabel and sort RGCs, followed by RNA sequencing (RNA-seq) (Baizabal et al., 2018; Hrvatin et al., 2014). We stained dissociated neocortical cells with PAX6 and TBR2 antibodies and sorted PAX6+ RGCs (Figure 6A). Compared to unsorted cell suspensions, sorted cell suspensions showed enrichment for PAX6+ RGCs (Figure 6B). We sequenced RNA libraries from five control and four *Yap;Taz* dKO biological replicates, with each replicate being obtained from two control embryos or three dKO embryos. Transcript counts confirmed high expression of RGC markers such as *Pax6* and *Nestin* and low expression of IPC and neuronal markers such as *Eomes*/*Tbr2* and *Tubb3*/*Tuj1* in sorted RGCs (Figure 6C). Consistent with YAP/TAZ being transcriptional coactivators, far more genes were downregulated than were upregulated in dKO RGCs when compared to control RGCs, with 336 genes being downregulated and 95 genes being upregulated with a fold change of ≥1.5 and an adjusted *P* value of ≤0.05 (Figure 6D, Supplementary Table 1). Gene set enrichment analysis using GO Biological Process gene sets revealed that, whereas no gene sets were significantly upregulated (with an FDR ≤0.25) in dKO RGCs, the top downregulated gene sets were Hippo signaling, Motile cilium assembly, and Cell substrate junction organization (Figure 6E, Supplementary Table 2). These gene expression changes correlate with and help explain our finding that YAP/TAZ are required for maintaining the proliferative capacity of RGCs, for generating ependymal cells, and for maintaining the structural organization of the neuroepithelium.

**Figure 6.**
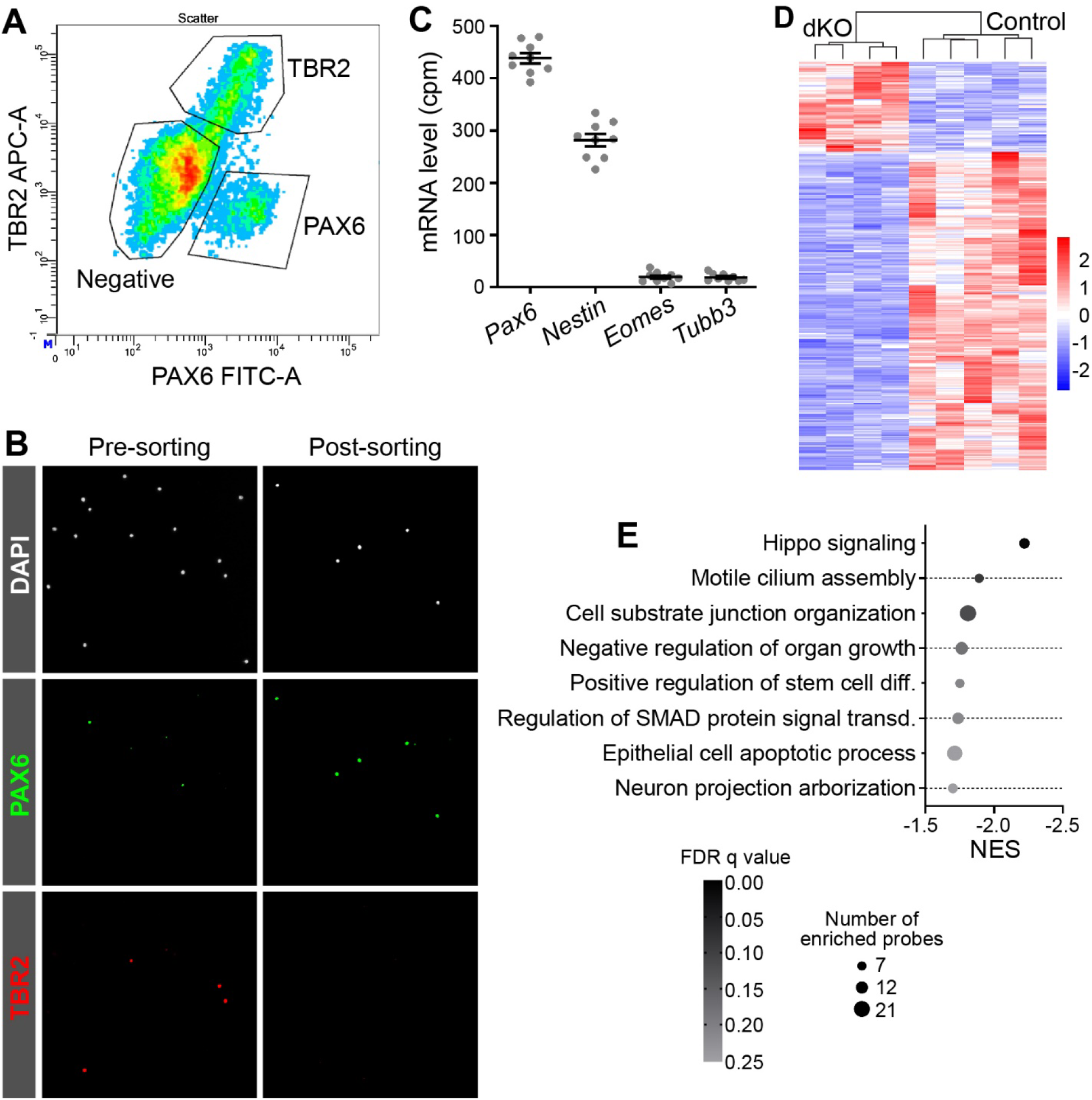
Transcriptional profiling of *Yap;Taz* dKO and control RGCs. (**A**) Representative plot showing sorting gates for PAX6+ RGCs and TBR2+ IPCs from E16.5 cortices. Double negative cells are cortical neurons. (**B**) Cell suspensions stained for DAPI, PAX6, and TBR2 before and after sorting of PAX6+ RGCs. (**C**) mRNA levels obtained from RNA-sequencing analysis of sorted E16.5 control and *Yap;Taz* dKO RGCs. (**D**) Heatmap of differentially expressed genes in E16.5 *Yap;Taz* dKO versus control RGCs with a fold change of ≥1.5 and an adjusted *P* value of ≤0.05. (**E**) Gene set enrichment analysis using the GO Biological Process gene sets, comparing the transcriptomes of *Yap;Taz* dKO versus control RGCs. Only enriched gene sets with an FDR q value of ≤0.25 are shown. For gene sets with similar probes, the gene set with the most enriched probes is shown.

## Discussion

By deleting both *Yap* and *Taz* in RGCs of the developing cortex, we have revealed that YAP/TAZ are required for maintaining the proliferative potential and structural organization of cortical RGCs and for the formation of the ependymal layer. YAP and TAZ function redundantly, and the loss of both results in hydrocephaly and reduced numbers of cortical projection neurons. Our RNA-seq analysis of sorted RGCs reveals the transcriptional basis for the functions of YAP/TAZ during cortical development.

YAP/TAZ are transcription factors and are typically believed to function by regulating gene expression. Interestingly, we found that the subcellular localization of YAP/TAZ undergoes dynamic changes during cortical development; it progresses from being largely cytoplasmic and highly enriched at the apical/ventricular surface of RGCs during early stages of cortical development to being enriched in the nucleus of RGCs at late stages of cortical development. This observation raises the question of whether the various functions of YAP/TAZ during cortical development are mediated by the cytoplasmic or the nuclear pool of YAP/TAZ proteins or whether, like β-catenin in the WNT pathway, YAP/TAZ exert different functions in the cytoplasm and the nucleus.

We found that the cell-cycle speed of *Yap;Taz* dKO RGCs was slower than that of control RGCs at E14.5, well before the neuroepithelial structural defects were first observed at E16.5, suggesting that the proliferation defect is not simply secondary to the structural defects, although it is possible that the structural defects contribute to the proliferation defect at late stages of cortical development. Therefore, the function of YAP/TAZ in maintaining the proliferative capacity of cortical RGCs is probably mediated by the nuclear—presumably transcriptional—activity of YAP/TAZ.

Whether the maintenance of RGC adherens junctions is mediated by nuclear or cytoplasmic YAP/TAZ is a more complicated question. In our analysis, although YAP/TAZ proteins were largely depleted from the ventricular surface as early as E12.5, junctional defects were not observed until E16.5, which coincided with the cytoplasm-to-nucleus translocation of YAP/TAZ during E14.5 to E16.5. Furthermore, our RNA-seq analysis revealed downregulation of genes involved in cell substrate junction organization in *Yap;Taz* dKO RGCs at E16.5, at which stage only a small fraction of the ventricular surface showed junctional defects in the dKO cortex. These observations suggest that the junctional defect is caused by the loss of the nuclear transcriptional function of YAP/TAZ. However, a previous study concludes that cytoplasmic, not nuclear, function of YAP is required for junctional integrity, based on the observations that injecting lysophosphatidic acid (LPA) into the developing cortex caused junctional disruption and promoted nuclear translocation of YAP and that the overexpression of wild-type (WT) and a phospho-mimetic YAP mutant (S112D), which presumably mimics the cytoplasmic YAP, but not of a phospho-defective YAP mutant (S112A), which mimics the transcriptionally active nuclear YAP, partially rescued the junctional defect (Park et al., 2016). However, the rescue of LPA-induced junctional defects by WT YAP and YAP-S112D was very limited. In fact, only the apical localization of N-cadherin was partially restored. No restoration of the full junction was reported, and RGCs and IPCs remained scattered. Furthermore, based on the transcriptional activity assay performed by these authors, the phospho-mimetic—supposedly cytoplasmic—YAP-S112D had transcriptional activity similar to that of WT YAP, suggesting that YAP-S112D is not completely cytoplasmic. Lastly, many studies have shown that activated YAP/TAZ promote the expression of genes involved in epithelial–mesenchymal transition (Aharonov et al., 2020; Diepenbruck et al., 2014; Lamar et al., 2012), a process that involves junction dissolution. Therefore, it is possible that the active form of YAP, YAP-S112A, did not restore adherens junctions because its overexpression caused a gain-of-function effect. Together, we argue that it remains to be determined whether YAP/TAZ act in the cytoplasm, in the nucleus, or in both compartments to maintain the structural organization of the neuroepithelium. Approaches, preferably genetic approaches, that enable specific perturbation of YAP/TAZ function in one compartment without causing secondary changes in YAP/TAZ activity in the other compartment, are needed to address this question.

In the ferret and human neocortices, YAP is necessary and sufficient for the proliferation of outer (or basal) RGCs (oRG) (Kostic et al., 2019), a type of progenitor cells that are abundant in the developing gyrencephalic neocortices but sparse in the developing lissencephalic mouse neocortex. Our finding that YAP/TAZ maintain the proliferative potential of apical RGCs in the mouse neocortex indicates that the role of YAP/TAZ in RGCs is evolutionarily conserved, whether in the apical RGCs of the lissencephalic mouse neocortex or in the oRG of gyrencephalic neocortices. In both types of neocortices, YAP/TAZ are not expressed in TBR2+ IPCs [this study and (Kostic et al., 2019)]. Whether YAP contributed to the expansion of the oRG population in gyrencephalic species over the course of evolution remains to be determined. One study showed that YAP/TAZ are required for the junctional integrity of the cerebellar neuroepithelium (Hughes et al., 2020), which is similar to our finding in the cortex. However, no defects in the proliferation capacity of cerebellar RGCs or granule neuron precursors were reported, although the *Yap;Taz* dKO cerebellum was smaller than the control cerebellum.

Although ependymal cells are born during mid-to-late embryonic development, ependymal cell differentiation and the formation of cilia occur much later, during the first postnatal weeks (Kyrousi et al., 2015; Spassky et al., 2005). YAP/TAZ depletion resulted in the complete loss of ependymal cells in P14 cortex, the earliest stage at which we could detect the definitive ependymal cell marker S100β. Whether this loss occurred because YAP is required for the specification of the ependymal cell fate or for the differentiation, maturation, or survival of ependymal cells or whether it was secondary to the reduced number of RGCs or the structural defects in the dKO cortex remains unknown. Addressing this question requires new tools that enable the specific deletion of *Yap;Taz* in the ependymal cell lineage or new markers that enable the detection of ependymal cells at embryonic stages. Even GemC1 and Mcidas, which are expressed in RGCs that will give rise to ependymal cells and are probably the earliest ependymal cell markers known, are not detectable in the cortex at embryonic stages (Kyrousi et al., 2015).

## Supporting information

Differentially expressed genes

Gene set enrichment analysis

## Acknowledgments

We thank Dr. Eric Olson for sharing the *Yap* and *Taz* floxed mice; Dr. Richard Ashmun and the St. Jude Flow Cytometry & Cell Sorting facility, Dr. Jennifer Peters and the St. Jude Cell & Tissue Imaging Center, The Hartwell Center for Biotechnology, Jim Houston and Kimberley Lowe in the Department of Developmental Neurobiology Flow Cytometry Core, and Maria Kinsey and Shanshan Kong in the Cao Lab for assistance with data acquisition and analysis; Dr. Abbas Shirinifard for implementing and customizing the cell quantification method with StarDist and OMERO; Drs. Corey Harwell and José-Manuel Baizabal at Harvard Medical School for sharing their MARIS protocol; and Dr. Keith A. Laycock for scientific editing of the manuscript.

---

**Supplementary Table 1. Differentially expressed genes in *Yap;Taz* dKO RGCs**

Differentially expressed genes in E16.5 *Yap;Taz* dKO RGCs compared to control RGCs with a fold change of ≥1.5 and an adjusted *P* value of ≤0.05.

**Supplementary Table 2. Enriched GO Biological Process gene sets in *Yap;Taz* dKO RGCs**

Gene set enrichment analysis comparing the transcriptomes of E16.5 *Yap;Taz* dKO versus control RGCs, showing gene sets with an FDR q value of ≤0.25.

## References

Aharonov, A., Shakked, A., Umansky, K.B., Savidor, A., Genzelinakh, A., Kain, D., Lendengolts, D., Revach, O.-Y., Morikawa, Y., Dong, J., Levin, Y., Geiger, B., Martin, J.F., Tzahor, E., 2020. ERBB2 drives YAP activation and EMT-like processes during cardiac regeneration. Nature Cell Biology 22, 1346–1356. 10.1038/s41556-020-00588-4

Allan, C., Burel, J.M., Moore, J., Blackburn, C., Linkert, M., Loynton, S., Macdonald, D., Moore, W.J., Neves, C., Patterson, A., Porter, M., Tarkowska, A., Loranger, B., Avondo, J., Lagerstedt, I., Lianas, L., Leo, S., Hands, K., Hay, R.T., Patwardhan, A., Best, C., Kleywegt, G.J., Zanetti, G., Swedlow, J.R., 2012. OMERO: flexible, model-driven data management for experimental biology. Nat Methods 9, 245–253. 10.1038/nmeth.1896

Anders, S., Pyl, P.T., Huber, W., 2015. HTSeq--a Python framework to work with high-throughput sequencing data. Bioinformatics 31, 166–169. 10.1093/bioinformatics/btu638

Baizabal, J.M., Mistry, M., García, M.T., Gómez, N., Olukoya, O., Tran, D., Johnson, M.B., Walsh, C.A., Harwell, C.C., 2018. The Epigenetic State of PRDM16-Regulated Enhancers in Radial Glia Controls Cortical Neuron Position. Neuron 99, 239–241. 10.1016/j.neuron.2018.06.031

Diepenbruck, M., Waldmeier, L., Ivanek, R., Berninger, P., Arnold, P., van Nimwegen, E., Christofori, G., 2014. Tead2 expression levels control the subcellular distribution of Yap and Taz, zyxin expression and epithelial-mesenchymal transition. J Cell Sci 127, 1523–1536. 10.1242/jcs.139865

Dobin, A., Davis, C.A., Schlesinger, F., Drenkow, J., Zaleski, C., Jha, S., Batut, P., Chaisson, M., Gingeras, T.R., 2013. STAR: ultrafast universal RNA-seq aligner. Bioinformatics 29, 15–21. 10.1093/bioinformatics/bts635

Eder, N., Roncaroli, F., Domart, M.-C., Horswell, S., Andreiuolo, F., Flynn, H.R., Lopes, A.T., Claxton, S., Kilday, J.-P., Collinson, L., Mao, J.-H., Pietsch, T., Thompson, B., Snijders, A.P., Ultanir, S.K., 2020. YAP1/TAZ drives ependymoma-like tumour formation in mice. Nature Communications 11, 2380. 10.1038/s41467-020-16167-y

Englund, C., Fink, A., Lau, C., Pham, D., Daza, R.A.M., Bulfone, A., Kowalczyk, T., Hevner, R.F., 2005. Pax6, Tbr2, and Tbr1 Are Expressed Sequentially by Radial Glia, Intermediate Progenitor Cells, and Postmitotic Neurons in Developing Neocortex. The Journal of Neuroscience 25, 247–251. 10.1523/jneurosci.2899-04.2005

Gorski, J.A., Talley, T., Qiu, M., Puelles, L., Rubenstein, J.L., Jones, K.R., 2002. Cortical excitatory neurons and glia, but not GABAergic neurons, are produced in the Emx1-expressing lineage. J Neurosci 22, 6309–6314.

Greig, L.C., Woodworth, M.B., Galazo, M.J., Padmanabhan, H., Macklis, J.D., 2013. Molecular logic of neocortical projection neuron specification, development and diversity. Nat Rev Neurosci 14, 755–769. 10.1038/nrn3586

Hrvatin, S., Deng, F., O’Donnell, C.W., Gifford, D.K., Melton, D.A., 2014. MARIS: method for analyzing RNA following intracellular sorting. PLoS One 9, e89459. 10.1371/journal.pone.0089459

Huang, Z., Hu, J., Pan, J., Wang, Y., Hu, G., Zhou, J., Mei, L., Xiong, W.-C., 2016. YAP stabilizes SMAD1 and promotes BMP2-induced neocortical astrocytic differentiation. Development (Cambridge, England) 143, 2398. 10.1242/dev.130658

Hughes, L.J., Park, R., Lee, M.J., Terry, B.K., Lee, D.J., Kim, H., Cho, S.-H., Kim, S., 2020. Yap/Taz are required for establishing the cerebellar radial glia scaffold and proper foliation. Developmental Biology 457, 150–162. https://doi.org/10.1016/j.ydbio.2019.10.002

Kostic, M., Paridaen, J.T.M.L., Long, K.R., Kalebic, N., Langen, B., Grübling, N., Wimberger, P., Kawasaki, H., Namba, T., Huttner, W.B., 2019. YAP Activity Is Necessary and Sufficient for Basal Progenitor Abundance and Proliferation in the Developing Neocortex. Cell Reports 27, 1103–1118.e1106. https://doi.org/10.1016/j.celrep.2019.03.091

Kriegstein, A., Alvarez-Buylla, A., 2009. The glial nature of embryonic and adult neural stem cells. Annu Rev Neurosci 32, 149–184. 10.1146/annurev.neuro.051508.135600

Kwan, K.Y., Sestan, N., Anton, E.S., 2012. Transcriptional co-regulation of neuronal migration and laminar identity in the neocortex. Development (Cambridge, England) 139, 1535–1546. 10.1242/dev.069963

Kyrousi, C., Arbi, M., Pilz, G.A., Pefani, D.E., Lalioti, M.E., Ninkovic, J., Götz, M., Lygerou, Z., Taraviras, S., 2015. Mcidas and GemC1 are key regulators for the generation of multiciliated ependymal cells in the adult neurogenic niche. Development (Cambridge, England) 142, 3661–3674. 10.1242/dev.126342

Lamar, J.M., Stern, P., Liu, H., Schindler, J.W., Jiang, Z.-G., Hynes, R.O., 2012. The Hippo pathway target, YAP, promotes metastasis through its TEAD-interaction domain. Proceedings of the National Academy of Sciences 109, E2441–E2450. 10.1073/pnas.1212021109

Lavado, A., He, Y., Pare, J., Neale, G., Olson, E.N., Giovannini, M., Cao, X., 2013. Tumor suppressor Nf2 limits expansion of the neural progenitor pool by inhibiting Yap/Taz transcriptional coactivators. Development (Cambridge, England) 140, 3323–3334. 10.1242/dev.096537

Lavado, A., Park, J.Y., Paré, J., Finkelstein, D., Pan, H., Xu, B., Fan, Y., Kumar, R.P., Neale, G., Kwak, Y.D., McKinnon, P.J., Johnson, R.L., Cao, X., 2018. The Hippo Pathway Prevents YAP/TAZ-Driven Hypertranscription and Controls Neural Progenitor Number. Developmental Cell 47, 576–591.e578. https://doi.org/10.1016/j.devcel.2018.09.021

Lavado, A., Ware, M., Pare, J., Cao, X., 2014. The tumor suppressor Nf2 regulates corpus callosum development by inhibiting the transcriptional coactivator Yap. Development (Cambridge, England) 141, 4182–4193. 10.1242/dev.111260

Law, C.W., Chen, Y., Shi, W., Smyth, G.K., 2014. voom: Precision weights unlock linear model analysis tools for RNA-seq read counts. Genome Biol. 15, R29. 10.1186/gb-2014-15-2-r29

Liu, W.A., Chen, S., Li, Z., Lee, C.H., Mirzaa, G., Dobyns, W.B., Ross, M.E., Zhang, J., Shi, S.-H., 2018. PARD3 dysfunction in conjunction with dynamic HIPPO signaling drives cortical enlargement with massive heterotopia. Genes & Development 32, 763–780. 10.1101/gad.313171.118

Ohata, S., Alvarez-Buylla, A., 2016. Planar Organization of Multiciliated Ependymal (E1) Cells in the Brain Ventricular Epithelium. Trends Neurosci 39, 543–551. 10.1016/j.tins.2016.05.004

Park, R., Moon, U.Y., Park, J.Y., Hughes, L.J., Johnson, R.L., Cho, S.-H., Kim, S., 2016. Yap is required for ependymal integrity and is suppressed in LPA-induced hydrocephalus. Nature Communications 7, 10329. 10.1038/ncomms10329

Pilaz, L.J., McMahon, J.J., Miller, E.E., Lennox, A.L., Suzuki, A., Salmon, E., Silver, D.L., 2016. Prolonged Mitosis of Neural Progenitors Alters Cell Fate in the Developing Brain. Neuron 89, 83–99. 10.1016/j.neuron.2015.12.007

Schmidt, U., Weigert, M., Broaddus, C., Myers, G., 2018. Cell Detection with Star-Convex Polygons, in: Frangi, A.F., Schnabel, J.A., Davatzikos, C., Alberola-López, C., Fichtinger, G. (Eds.), Medical Image Computing and Computer Assisted Intervention – MICCAI 2018. Springer International Publishing, Cham, pp. 265–273, https://doi.org/10.1007/978-3-030-00934-2_30

Shao, W., Yang, J., He, M., Yu, X.-Y., Lee, C.H., Yang, Z., Joyner, A.L., Anderson, K.V., Zhang, J., Tsou, M.-F.B., Shi, H., Shi, S.-H., 2020. Centrosome anchoring regulates progenitor properties and cortical formation. Nature 580, 106–112. 10.1038/s41586-020-2139-6

Spassky, N., Merkle, F.T., Flames, N., Tramontin, A.D., García-Verdugo, J.M., Alvarez-Buylla, A., 2005. Adult Ependymal Cells Are Postmitotic and Are Derived from Radial Glial Cells during Embryogenesis. The Journal of Neuroscience 25, 10. 10.1523/JNEUROSCI.1108-04.2005

Subramanian, A., Tamayo, P., Mootha, V.K., Mukherjee, S., Ebert, B.L., Gillette, M.A., Paulovich, A., Pomeroy, S.L., Golub, T.R., Lander, E.S., Mesirov, J.P., 2005. Gene set enrichment analysis: A knowledge-based approach for interpreting genome-wide expression profiles. Proceedings of the National Academy of Sciences 102, 15545. 10.1073/pnas.0506580102

Taverna, E., Götz, M., Huttner, W.B., 2014. The Cell Biology of Neurogenesis: Toward an Understanding of the Development and Evolution of the Neocortex. Annual Review of Cell and Developmental Biology 30, 465–502. 10.1146/annurev-cellbio-101011-155801

Tronche, F., Kellendonk, C., Kretz, O., Gass, P., Anlag, K., Orban, P.C., Bock, R., Klein, R., Schutz, G., 1999. Disruption of the glucocorticoid receptor gene in the nervous system results in reduced anxiety. Nat Genet 23, 99–103.

Villalba, A., Götz, M., Borrell, V., 2021. Chapter One - The regulation of cortical neurogenesis, in: Bashaw, G.J. (Ed.), Current Topics in Developmental Biology. Academic Press, pp. 1–66. https://doi.org/10.1016/bs.ctdb.2020.10.003

Xin, M., Kim, Y., Sutherland, L.B., Qi, X., McAnally, J., Schwartz, R.J., Richardson, J.A., Bassel-Duby, R., Olson, E.N., 2011. Regulation of insulin-like growth factor signaling by Yap governs cardiomyocyte proliferation and embryonic heart size. Sci Signal 4, ra70.

Zheng, Y., Pan, D., 2019. The Hippo Signaling Pathway in Development and Disease. Developmental Cell 50, 264–282. 10.1016/j.devcel.2019.06.003

